# Xist IncRNA forms silencing granules that induce heterochromatin formation and repressive complexes recruitment by phase separation

**DOI:** 10.1101/351015

**Authors:** Andrea Cerase, Alexandros Armaos, Fernando Cid, Philip Avner, Gian Gaetano Tartaglia

**Affiliations:** EMBL-Rome, Via Ramarini 32, 00015 Monterotondo (RM), Italy; Centre for Genomic Regulation (CRG), The Barcelona Institute of Science and Technology, Dr Aiguader 88, 08003 Barcelona, Spain; Universitat Pompeu Fabra (UPF), 08003 Barcelona, Spain; Institucio Catalana de Recerca i Estudis Avançats (ICREA), 23 Passeig Lluis Companys, 08010 Barcelona, Spain; Equal contribution

**Keywords:** Xist lncRNA, long non-coding RNAs (lncRNA), tandem repeats, epigenetics, X chromosome inactivation (XCI), RNA-granules, stress granules, paraspeckles, liquid-phase separation, heterochromatin, intrinsically disordered proteins, intrinsically disordered RNAs, high-order chromatin structure, Polycomb Repressive Complexes 1/2 (PRC1/2), nuclear matrix, RNA-protein interaction, RNA secondary structure, ribonucleoprotein (RBN)

## Main text

Long non-coding RNAs (lncRNAs) are RNA molecules longer than 200 bases that lack coding potential^1,2^. They represent a significant portion of the cell transcriptome^3^ and work as activators or repressors of gene transcription acting on different regulatory mechanisms^4–6^. Indeed, lncRNAs can act as macro-scaffolds for protein recruitment^7–14^ and behave as guides and sponges for titrating RNA and proteins, influencing transcription at regulatory regions or triggering transcriptional interference^15–17^.

The ability of RNA to scaffold protein interactions has been shown to contribute to the formation of membrane-less organelles such as paraspeckles^18^ and stress granules^19^. These assemblies, mainly composed of RNAs and proteins, aggregate through a process of liquid-liquid phase separation^18–20^. Formation of ribonucleoprotein granules is an evolutionary conserved mechanism for cells to respond to environmental changes^61^ and favors the confinement of enzymes and nucleic acids in discrete regions of the nucleus or cytoplasm^21^. Structurally disordered and nucleic acid binding domains promote protein-protein and protein-RNA interactions in the granules^22^. Especially intrinsically disordered proteins, which are enriched in polar amino acids such as arginine, glycine and phenylalanine, have been shown to promote phase transitions in the cell^23^.

Recent experiments unveiled an interesting link between the process of phase separation and heterochromatinization^24–26^. Indeed, several lncRNAs such as HotAir^27^, ARNIL^28^, Airn^29^ and Xist^4^ lncRNAs can induce heterochromatin formation to different extent (reviewed in Long et al.^6^). Xist in particular, the master regulator of X chromosome inactivation, has been shown to induce large-scale heterochromatinization of the entire X-chromosome^4^ accumulating in large granule-like assemblies that can be investigated by super-resolution microscopy^30,31^ (**Fig. 1A**). Female cells show ~100-150 Xist-containing complexes^30,31^ that largely resembles paraspeckles and stress granule in size and protein composition (see below). Intriguingly, Xist assemblies measure around 100-200 nm in diameter^31^, comparable with stress granules and paraspeckles dimensions (~100- 300 nm and ~200-500 nm in diameter, respectively^32,19^) (**Fig. 1A**). We note that the slight discrepancy in size of ribonucleoprotein assemblies is due to the diverse functions and compositions^4,19,32^. Other lncRNAs such as HotAir, Malat1 or Airn seem to form smaller *puncti* assemblies, rather than granule-like structures^33,34^. However, a stringent comparative analysis of these lncRNAs and their interacting proteins, by means of super-resolution microscopy is still missing, hampering a more formal comparison.

**Fig 1.**
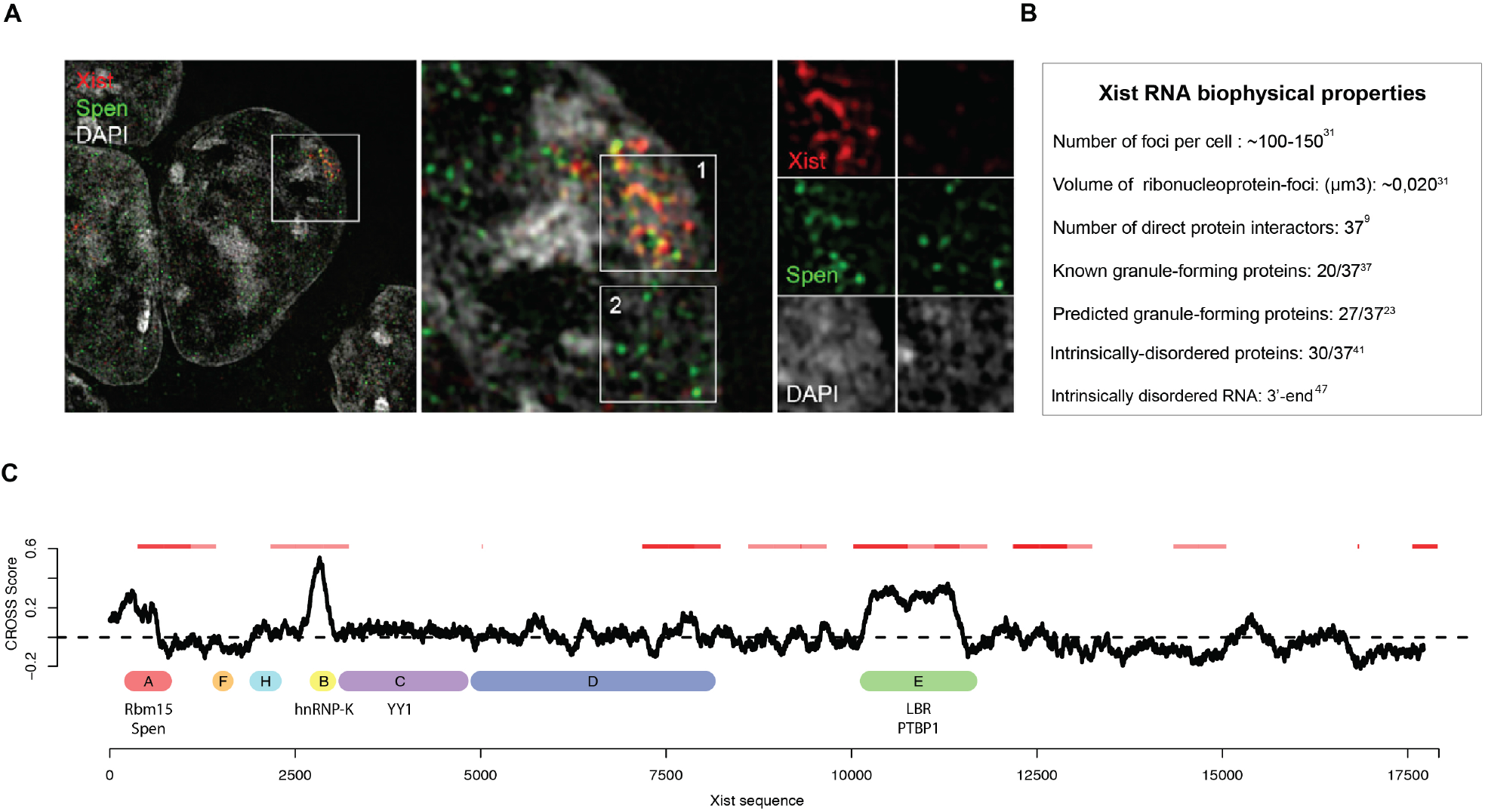
**A)** Xist forms silencing granules and is enriched for granule binding proteins. Left, representative image of Xist silencing granules^7^. With permission from the editor, *Cell Report*, volume 12, pages 562-72, Cell-Press, 2015 (Open Access, Creative Commons Attribution License (CC BY). **B)** Quantification of Xist-granule bio-physical properties and cellular distribution. Xist biophysical properties are shown^31^. **C)** Xist secondary structure prediction (*y* axis: CROSS score^47^; *x* axis: Xist sequence mouse mm10 (MGI:98974). Below CLIP-validated Xist binding partners^10,11,13^. Above binding sites regions of granule-forming interactions ^18–20,32^

Supporting our observation that Xist assemblies are granule-like, we report that Xist interactions involve a copious number of proteins that phase separate. Indeed, Xist physically binds to a few dozen partners, although more than 600 proteins have been found to associate through non-direct, yet potentially biologically-relevant interactions (“putative” Xist network, see Materials and Methods for details)^9^. Using literature^16,19,20,32^, computational and experimental data, we found that the Xist direct interactome contains proteins that are prone to form granules. More specifically, the proteins binding to Xist show significant overlap with paraspeckles (29 out of 37 Xist binders are Neat1 interactors too; p-value < 0.002, Fisher’s test computed using 94 eCLIP experiments; 2 of the interactions (Rmb14 and Tardb1) are among the 7 paraspeckle architectural binders^18,35,36^; See **Table 1**) and stress-granule composition (20 out 37 direct interactors, with enrichment p-value < 0.0001 and 58 out of 631 enrichment p-value < 0.0002 considering all associations ^7–12^; Fisher’s test enrichments calculated using 240 stress-granule core components^37^; see **Figure 1B** and online **Table S1,** https://tinyurl.com/y9ygtyn4). Moreover, all the 37 direct interactors are predicted to form large ribonucleoprotein assemblies (granule propensity scores > 0) and 27/37 are significantly prone to phase separate (granule propensity scores > 1). Furthermore, 24% of the putative Xist interactome is also enriched of granule-forming proteins (146 proteins; for this analysis we used the catGRANULE method^23^; all values are available at https://goo.gl/8CBwMp) and https://tinyurl.com/yd3tjzs5); **Fig. 1B**). Among the Xist interactions, Rbm14 and Tardbp1 are reported to be critical in paraspeckles formation^18^ while Hnrnp proteins, key components of stress-granules, are necessary for Xist silencing (HnrnpU)^38^ and Polycomb recruitment (HnrnpK) activities, respectively^39^. By comparison, we analysed the putative interactome of other lncRNAs such as HotAir and Malat1 (also known as NEAT2)^40^. We found a poor overlap with the interactomes of paraspeckles and stress-granules (HotAir: 0 out of 6 are Neat1 interactors, p-value=1 and 0 out of 6 are stress-granules components, p-value=1; Malat1: 5 out of 26 are Neat1 interactors, p-value=0.6, and 7 out of 26 are stress-granules components; p-value=0.2, Fisher’s test). HotAir and Neat1 interactomes are significantly smaller than Xist’s, supporting the observation that these RNAs form *puncti* assemblies^33,34^. These observations may indicate that not all lncRNAs are enriched in granule-forming proteins, possibly depending on RNA length, spatial conformation and protein interactions.

**Table1.**
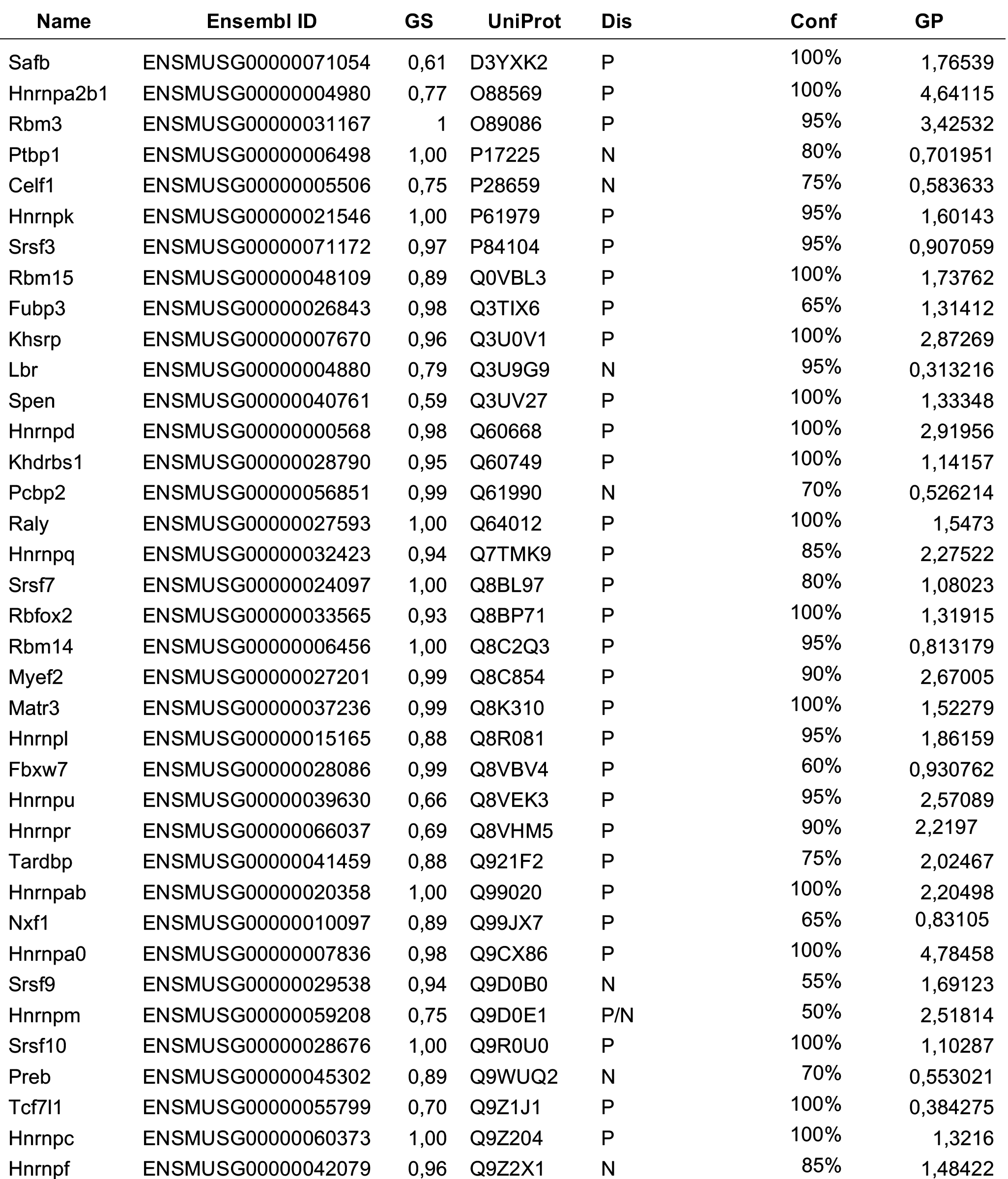
Xist direct interactome enrichment for granule forming proteins. Tables also shown intrinsically disordered proteins (Dis), the degree of confidence (Conf) and the granule propensity (GP).

Structural disorder and nucleic acid-binding are properties of proteins that assemble into ribonucleoprotein (RNP) granules^23^. We observe a strong enrichment of structurally disordered proteins in the Xist interactome (18 out of 37, or 50%, physical interactions are predicted to be highly disordered with p-value enrichment < 0.0001; 184 out of 631, or 30%, total Xist associations are disordered, p=0.005; 111 out of 631 disordered-prone proteins are also forming stress granules, p-value = 0.001; calculations of structural disorder have been carried out using *clever*Machine, values are available at https://goo.gl/eyX2bq)^41^; p-values were computed with *Chi-square* with Yates’ correction; **Fig. 1C**). These results are truly remarkable, since the fraction of proteins predicted to be structurally disordered in *M. musculus* is 22% (i.e., 3467 out 15338 non-redundant; Uniprot database). Among the disordered-prone proteins, HnrnpA2B1 and HnrnpQ, Ptbp1, Ttcf7l1 and Spen show high propensity for granule formation and are core constituents of paraspeckles^18^ and stress-granules^19,20^. We stress that the presence of structurally disordered proteins is important for the process of phase separation, as showed by different experimental and computational studies^22,24,25^. Indeed, eukaryotic proteomes contain intrinsically unfolded and repetitive regions in granule-forming proteins^23^, which confer them an intrinsic ability to promote protein-protein and protein-nucleic-acid interactions^42,43,44^. As a comparison, we analyzed the content of structurally disordered proteins in the interactomes of HotAir and Malat1, but the enrichments are not significant (highly disordered proteins: 2 out of 6 for HotAir, with p-value=0.7 and 6 out of 26 for Malat1 with p-value=0.4), indicating a substantial difference with Xist RNA interaction network.

In the large spectrum of activities, RNA structure plays a central role and dictates precise functionalities by creating spatial patterns and alternative conformations and binding sites for proteins^45^. As for Xist RNA, six conserved repetitive regions (tandem repeats), named A to F, have been reported to be essential for its function^45^ (**Fig. 1C**). Xist repeats are conserved in mammalian vertebrates but considerable variation in the copy number is observed, with the exception of the A repeat region, which is conserved both in terms of copy number and consensus sequence^45^. In agreement with dimethyl sulfate (DMS)-sensitivity experiments^46^ and predictions of RNA structure^47^, repeats A, B and E are highly structured (CROSS predictions are available at https://goo.gl/yzqUjS). While repeats A and B are conserved across species and show a high degree of structural content, the 3’ region of Xist is variable and predicted to largely be singlestranded^47^. RNA-binding predictions of Xist interactions^9,47^ indicate that Rep D and E and A (and B to a lesser extent) are crucial for Xist interactions with granule-forming proteins (**Fig. 1C**). This result is in line with the recent observation that structured regions attract granule-forming proteins^48,61^.

Could the formation granule-like assembly be relevant for Xist function? In a recent publication, we investigated the interaction of Xist with Ezh2 and other proteins of the PRC2 complex^49^. Using randomized repeat A as a control, we estimated Ezh2 to bind Xist with high affinity but low specificity^49^. Our findings are in good agreement with 3D-SIM data, showing poor overlap between Xist and PRC2^30^. Yet, data by Stochastic Optical Reconstruction Microscopy (STORM) indicate non-random association of Xist and PRC2^50^. Thus, Xist and PRC2 are closer than expected by chance, but their interactions are unstable, similarly to those established by disordered proteins within stress granules. This observation is particularly pertinent if we consider that ribonucleoprotein granules are in a dynamic intermediate metastable state^51^ that makes their purification particularly difficult. Indeed, the fast exchange with the *cell milieu* impedes the isolation of the liquid phase assemblies^52,53^.

At present, the interaction between Xist RNA and certain subunits of the PRC2 complex RNA and the molecular mechanisms of recruitment remain controversial. Some literature reports evidence arguing in favor^54,55^ while other is against^30,39,56^. While the Xist-PRC2 interaction *in vitro* is strong, it is possible that the interaction of the PRC2 to repeat A *in vivo* has such a fast kinetics that prevents it to be captured by most studies^50^. The main point of discussion is the observation of a strong interaction between repeat A and PRC2 *in vitro*^54,57^, which seems to be dispensable *in vivo*^39,56^ as a form of Xist lacking the repeat A^58^, can still induce *de novo* recruitment of PRC2 (and similarly of PRC1. These observations may indicate that more than one Xist region may be involved in this process. Indeed recent work has shown that the *de novo* recruitment of PRC2 is mediated by Xist RepB and non-canonical PRC1 variants^28,37,56^.

Here we propose that Xist exerts its functions through the formation of silencing granules in which repressive complexes are recruited by phase separation. More precisely, we suggest that non-canonical recruitment of repressive complexes PRC1 is promoted or reinforced by the formation of phase-separated large assemblies. In this scenario, the primary *de novo* PRC1 recruitment is mediated by Xist RepB^39^ and PRC1-mediated H2A ubiquitination may trigger PRC2 recruitment as previously shown^39,56,59,60^ (**Fig. 2**). PRC1 recruitment may be further strengthened via the interaction with other intrinsically disordered domain-containing proteins binding to Xist, thus mediating further recruitment and oligomerization (see **Fig.2**). It is possible that other disorder-containing domains proteins such as Saf-A (repeat A) or Matr3 and Ciz1 (repeat E) may mediate the PRC1 recruitment in the Xist body and trigger phase-separation (see **Table 1** **and** **Fig. 2**; data available at https://goo.gl/2o7L43) and https://goo.gl/hsKq2R). However, we cannot exclude that these interactions are mediated by other intrinsically-disordered proteins yet to be discovered binding the A-, D-3’end and the E-repeats (see **Fig. 2**). This multimerization driven by phase-separation may, in turn, trigger RNA Polymerase II (PolII) and basic transcription factors eviction, inducing gene-silencing and heterochromatinization. Supporting this hypothesis, we previously reported that i) Ring1b is relatively closer to Xist than other PRC1/2 components, as shown by super-resolution analysis, suggesting a non-chromatin mediated yet indirect contact^30^; ii) structurally-disordered and granule-forming PRC1 core components such as Rybp and Rnf2 (Ring1b) are highly prone to interact with Xist^9^, directly or indirectly, in agreement with experimental screenings^10,11,13^; iii) nuclear matrix proteins are highly prone to interact with Xist RNA via Xist E-repeat^9^, iv) Xist RNA has all the critical characteristic of an RNA-protein granule (see above).

**Fig 2.**
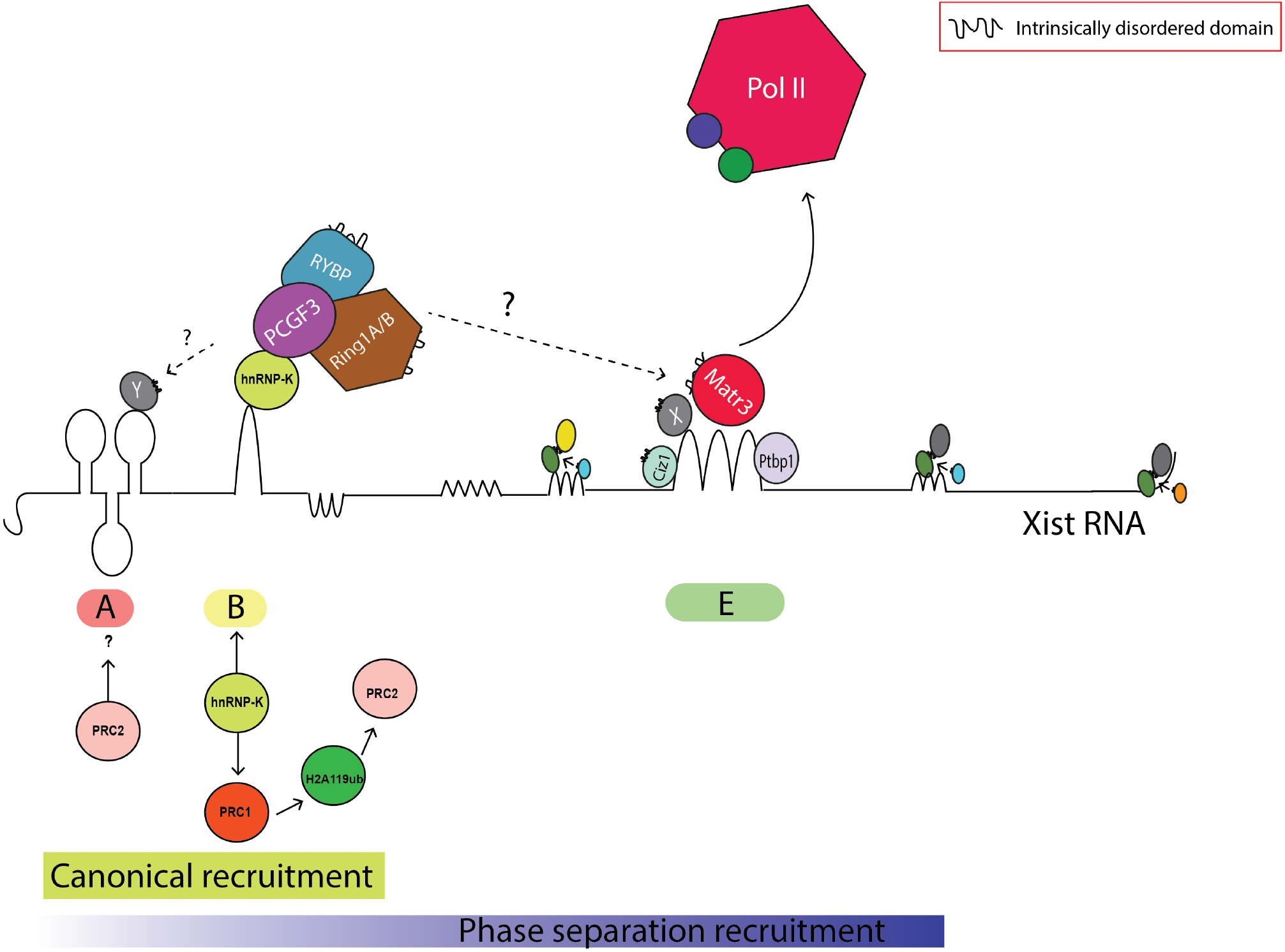
Phase separation and Xist RNA recruitment of repressive complexes. In addition to the canonical PRC2 recruitment that involve repeat B (indirect, PRC1-mediated recruitment), we suggest that Xist recruits PRC1 once in a phase-separated state through mediation of structurally disordered proteins binding repeat E and repeat A. PRC1 complex recruitment could occur through a direct interaction with repeat E (e.g., Ring1b, Rybp) or via Ciz1 and Matr3 (as well as other unknown proteins X, binding to repeat E and containing intrinsically disordered regions). It is possible that PRC1 interacts with repeat A in a phase-separated state after the initial *de novo* seed has been placed via an unknown disordered protein Y. H2A ubiquitination by PRC1 may induce PRC2 recruitment on the Xi as previously shown (see main text). We suggest that the oligomerization can further recruit repressive proteins and/or disordered proteins, evicting Pol IIand basic transcription factors, recruiting more structurally disordered proteins and in turn, inducing further granule formation, heterochromatinization and gene-repression.

In brief, we suggest that regardless of the type of interaction, which could be direct or indirect, components of the PRC1/2 repressive complexes are recruited into Xist granules through a mechanism of phase separation. Noticeably, a similar mechanism for the recruitment of protein complexes may be exploited by other lncRNAs as well and may not be limited to Xist RNA. This mechanism of action can be used by lncRNAs to recruit other repressive complexes to the heterochromatin and not being limited to PRC1/2 complexes. As our hypothesis is supported by correlative evidence, more experimental work has to be done in order to fully validate it.

## Acknowledgements

We would like to thank all members of the Avner and the Tartaglia groups, Greta Pintacuda, Mitchell Guttman and Kathrin Plath for critical reading of the manuscript. AC and PA were funded by an EMBL grant to PA (50800). The research leading to these results has been supported by European Research Council (RIBOMYLOME_309545), Spanish Ministry of Economy and Competitiveness (BFU2014-55054-P and BFU2017-86970-P) and “Fundació La Marató de TV3” (PI043296).

## Material and methods (in brief)

### Microscopy

Super resolution ImmunoRNA-FISH and Immuno-Fluorescence (IF) microscopy and analysis has been performed in previously-published research^30,31^.

### RNA Structure

We predicted the secondary structure of granule and non-granule transcripts using CROSS (Computational Recognition of Secondary Structure^47^. CROSS was developed to perform high-throughput RNA profiling. The algorithm predicts the structural profile (single-and double-stranded state) at single-nucleotide resolution using sequence information only and without sequence length restrictions (scores > 0 indicate double stranded regions).

### Protein-RNA interactions

*cat*RAPID *Global Score* was used to compute the interaction propensity of *Xist* with proteins^9^. Exploiting sequence information, the algorithm integrates local properties of protein and RNA structures into an overall binding propensity (scores > 0.5 indicate strong interactions). All the proteins analysed in this article were previously reported in our previous publication^9^ (including Xist putative interactome consisting of 631 proteins). We want to stress that the putative Xist interactome has a higher degree of false discovery rate (FDR)^7,8,10,11,13^, and therefore is less reliable than the putative direct interactome^9^.

### Granule propensity

Structural disorder, nucleic acid binding propensity and amino acid patterns such as arginine-glycine and phenylalanine-glycine are key features of proteins coalescing in granules^23^. These features were combined in a computational approach, *cat*GRANULE, that we employed to identify RBPs assembling into granules (scores > 0 indicate granule propensity).

